# Skeletal Muscle Stem Cell-Derived Myonuclei Adopt Divergent Terminal Transcriptional States in Adult and Aged Muscle In Response to a Hypertrophic Stimulus

**DOI:** 10.64898/2026.08.26.747125

**Authors:** Nicholas T. Thomas, Jensen Z. Goh, Kevin A. Murach, Christopher S. Fry, Charlotte A. Peterson, Ahmed Ismaeel, John J. McCarthy, Yuan Wen

**Affiliations:** Department of Physiology, College of Medicine, University of Kentucky, Lexington, KY, USA; Center for Muscle Biology, University of Kentucky, Lexington, KY, USA; Division of Biomedical Informatics, Department of Internal Medicine, Lexington, KY, USA; Department of Nutrition and Exercise Physiology, Department of Medical Pharmacology and Physiology, NextGen Precision Health Initiative, University of Missouri, Columbia, MO, USA; Department of Athletic Training and Clinical Nutrition, College of Health Science, University of Kentucky, Lexington, KY, USA; Department of Anatomy, Physiology, & Pharmacology, College of Veterinary Medicine, Auburn University, Auburn, AL, USA

**Author notes:** Co-Senior/Co-Corresponding Authors: Yuan Wen, MD/PhD, University of Kentucky, Lexington, KY 40536, John J. McCarthy, PhD, University of Kentucky, Lexington, KY 40536, Ahmed Ismaeel, PhD, Auburn University, Lexington, KY 40536, Auburn, AL, USA 36849.

**Keywords:** Stem cells, Aging, Exercise, Myonuclei

## Abstract

Skeletal muscle stem cells (MuSCs) give rise to a fusogenic cell population that provide new myonuclei to muscle fibers. Myonuclear functional heterogeneity has recently become appreciated, but the terminal identity of MuSC-Derived myonuclei remains undefined. We performed single-nucleus RNA-sequencing of myonuclei in Adult and Aged muscle to define MuSC-Derived and resident myonuclear responses to mechanical overload (MOV), which induces a hypertrophic stimulus. We found a MuSC-dependent induction of a youthful transcriptional signature in resident myonuclei after MOV in Aged muscle. Age determined terminal transcriptional states of MuSC-Derived myonuclei toward MTJ in Adult, NMJ in Aged, and muscle spindles in both ages. Microtubule-remodeling genes, *Macf1*, *Map1b*, and *Nav3*, along with the transcription factor *Runx1*, identified this post-fusion specialization with greater expression of these genes in Adult than in Aged MuSC-Derived myonuclei. In-silico transcription factor KO screen identified Runx1 as a regulator of post-fusion specialization and Esrrg as a driver of spindle (intrafusal) MuSC-Derived myonuclear maturation. By defining the age-associated fate of MuSC fusion to muscle fibers, we provide potential targets for modulating muscle plasticity.

**Highlights:**

- MuSC presence creates a more youthful myonuclear transcriptional signature in Aged muscle with mechanical overload (MOV).
- MOV-Responsive myonuclei depend on MuSCs in Aged but not Adult skeletal muscle
- MuSC-Derived myonuclei transcriptionally specialize to support the myotendinous junction in Adult muscle, the neuromuscular junction in Aged muscle, and muscle spindle (intrafusal) fibers in both ages.
- Pseudotime predicts *Runx1* may function as a master regulator of MuSC-Derived myonuclear specialization in response to MOV.

## Introduction

Skeletal muscle possesses remarkable capacity for adaptive remodeling in response to mechanical load that occurs with resistance exercise training. Such adaptive remodeling is essential for maintaining strength, metabolic health, and functional independence throughout the lifespan. With advancing age, impaired skeletal muscle plasticity contributes to sarcopenia and its clinical consequences including frailty, falls, fractures, loss of independence, and increased mortality. Although resistance exercise remains the most effective intervention, its benefits are limited by anabolic resistance, a poorly understood blunting of the adaptive response to mechanical loading.^1–3^

Skeletal muscle stem cells (MuSCs) are activated, proliferate, differentiate, and fuse into existing myofibers during mechanical loading, thereby contributing new myonuclei to the syncytium^4,5^. This myonuclear accretion accompanies hypertrophic adaptation and is generally reduced in aged muscle^6,7^. The skeletal muscle field has focused predominantly on the events preceding fusion (MuSC activation, proliferation, and differentiation) almost exclusively in the context of regeneration while a more fundamental question remains unanswered: what is the transcriptional fate of MuSC-Derived myonuclei once incorporated into a pre-existing myofiber during hypertrophy? Further, do MuSC-Derived myonuclei impact the transcriptome of existing resident myonuclei? The technical challenge of distinguishing newly incorporated MuSC-derived myonuclei from resident myonuclei has prevented understanding their functional contribution to adaptation.

This knowledge gap has profound implications for understanding age-related impairments in muscle plasticity. If MuSC-Derived myonuclei adopt specialized transcriptional programs that support hypertrophy, distinct from resident myonuclei, then age-related defects could arise at multiple levels; not only reduced fusion, but altered transcriptional programming of newly incorporated nuclei themselves. Conversely, resident myonuclei may possess intrinsic transcriptional deficits in aged muscle that limit adaptive capacity independent of MuSC contribution^8^. Disentangling these possibilities requires an experimental approach capable of resolving the transcriptional identities of MuSC-Derived versus resident myonuclei during the adaptive response to mechanical overload (MOV) induced by synergist ablation.

We designed the present study to address three questions: (1) What is the transcriptional fate of MuSC-Derived myonuclei during MOV? (2) Is the fate of MuSC-derived myonuclei altered with old age? (3) Do resident myonuclei in aged muscle show intrinsic transcriptional deficits in response to MOV, independent of MuSC contribution? To answer these questions, we generated a novel transgenic mouse model that allows inducible depletion of MuSCs (Pax7^CreER/+^:Rosa26^DTA/+^) combined with inducible recombination-independent myonuclear fluorescent labeling (HSA^rtTA^:TRE^H2B-GFP^) followed by single nucleus RNA-seq (snRNA-seq) of purified myonuclei. By comparing MuSC-retained and MuSC-depleted conditions across ages and loading states, we computationally identified MuSC-dependent transcriptional populations and dissected the independent effects of age on both newly incorporated and resident myonuclear transcriptomes.

## Results

### Myonuclear transcriptional profiles are responsive to aging, MOV, and MuSC depletion

Adult and Aged Pax7^CreER/+^-Rosa26^DTA/+^: HSA^rtTA^-TRE^H2B-GFP^ (Pax7-DTA:HSA-GFP) mice were first administered tamoxifen to deplete skeletal muscle stem cells (MuSC**^-^**) or vehicle to retain muscle stem cells (MuSC**^+^**), then following a two-week washout period, underwent synergist ablation surgery to induce MOV of the plantaris muscle or Sham surgery for 14 days. Doxycycline was provided via drinking water for the last 3 days of MOV to GFP-label myonuclei (Figure 1A). This experimental design resulted in 8 groups: Adult Sham MuSC**^+^**, Adult MOV MuSC**^+^**, Adult Sham MuSC**^-^**, Adult MOV MuSC**^-^**, Aged Sham MuSC**^+^**, Aged MOV MuSC**^+^**, Aged Sham MuSC**^-^**, and Aged MOV MuSC**^-^**. For each group, myonuclei were sorted from pooled plantaris muscles (n=2-3) using Fluorescence Activated Nuclear Sorting (FANS) for snRNA-seq analysis (Figure 1A). Myonuclear clusters were identified on the basis of myosin heavy chain (*Myh*) gene expression of known fiber-types of the plantaris muscle (*Myh1*, Type IIx; *Myh2*, Type IIa; *Myh4*, Type IIb) and the expression of key markers defining specialized myonuclear clusters including neuromuscular junction (NMJ, *Colq*), myotendinous junction (MTJ, *Col22a1*), muscle intrasfusal spindle (*Myh7*, *Myh7b*, *Myh6*, *Myl2*), and denervated (*Runx1*, *Gadd45a*, *Fbxo32*) myonuclei (Figure 1A).

**Figure 1:**
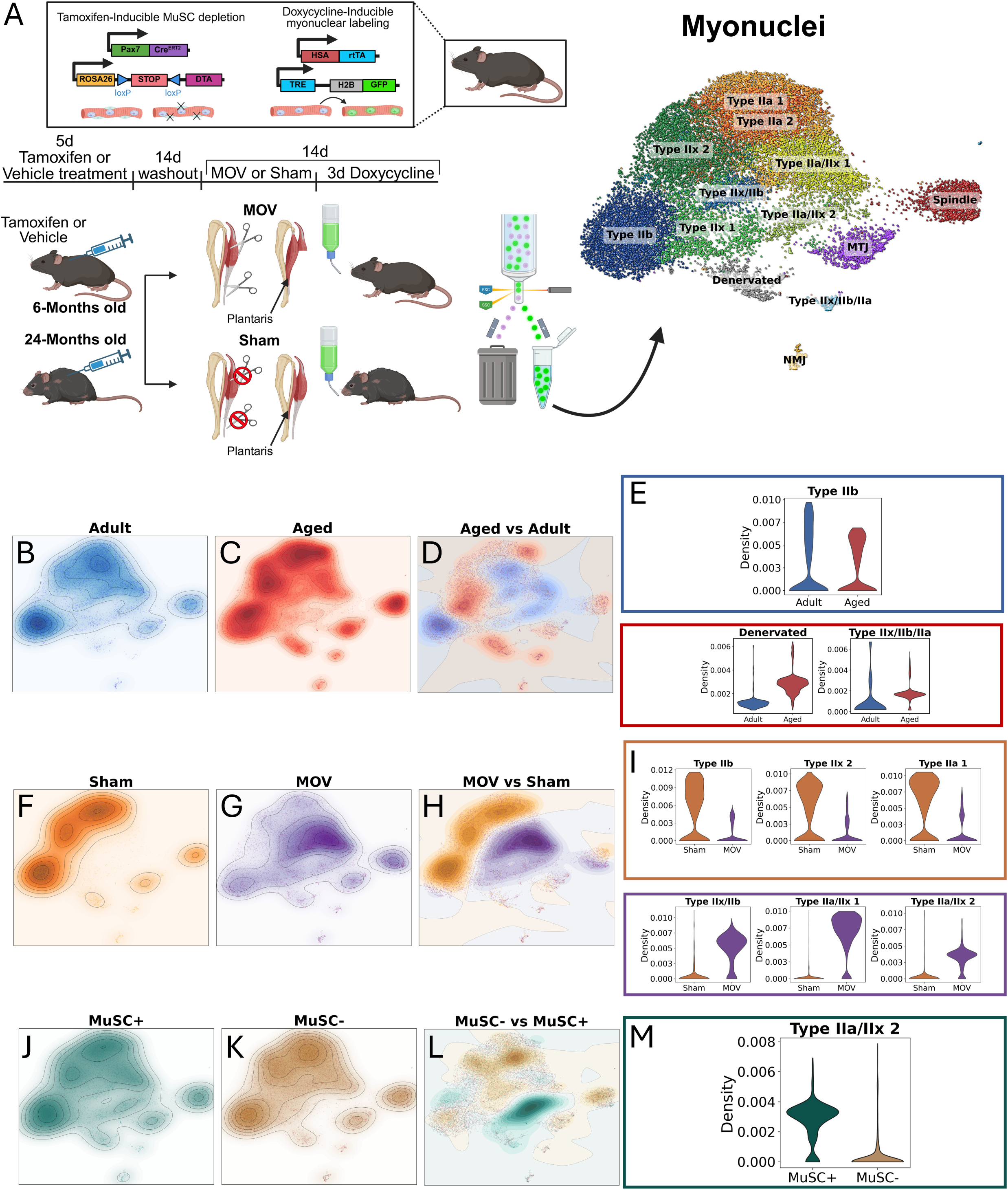
MOV, MuSC depletion, and aging affect distribution of myonuclear transcriptional profiles. A. Study design schematic showing genetic strategy, tamoxifen, doxycycline, and mechanical overload timing in Adult and Aged mice prior to fluorescent sorting and single nucleus RNA sequencing of myonuclei alongside uniform manifold approximation projection (UMAP) plot of myonuclei and myonuclear cluster identities, created in Biorender. B. Count-normalized density of myonuclear abundance within the integrated UMAP from Adult and C. Aged samples. D. Differential normalized density between Adult and Aged samples. E. Distribution of normalized density for Type IIb, Denervated, and Type IIx/IIb/IIa clusters per Adult and Aged conditions. F. Count-normalized density of myonuclear abundance within the integrated UMAP from Sham and G. MOV samples. H. Differential normalized density between Sham and MOV samples. I. Distribution of normalized density for Type IIb, Type IIx2, Type IIa 1, Type IIx/IIb, Type IIa/IIx 1, and Type IIa/IIx 2 myonuclear clusters per Sham and MOV conditions. J. Count-normalized density of myonuclear abundance within the integrated UMAP from MuSC**^+^** and K. MuSC**^-^** samples. L. Differential normalized density between MuSC**^+^** and MuSC**^-^** samples. M. Distribution of normalized density for the Type IIa/IIx 2 cluster per MuSC**^+^** and MuSC**^-^** conditions.

Shifts in myonuclear density within a uniform manifold approximation projection (UMAP) allowed interrogation of differences in myonuclear gene expression with each independent variable (all Adult vs all Aged samples, Figure 1B-E; all Sham vs all MOV samples, Figure 1F-I; all MuSC**^+^** vs all MuSC**^-^** samples, Figure 1J-M). High differential density of myonuclei within a region of the UMAP space defines condition-specific myonuclear transcriptional signatures. Differential density showed Aging primarily lowered the density of pure fast, glycolytic Type IIb myonuclei (Figure 1B-E) consistent with prior observations of lower Type IIb fiber frequency in plantaris muscle with advanced age^7^. Aging also induced an increase in the abundance of Denervated and Type IIx/IIb/IIa myonuclear clusters (Figure 1B-E). MOV induced the greatest overall shift in density compared to the Sham condition (Figure 1F-I), including a notable increase in the density of fast hybrid myonuclear clusters, Type IIx/IIb, Type IIa/IIx-1, and Type IIa/IIx-2 (Figure 1F-I). MuSC depletion had a relatively modest effect on myonuclear density across clusters, except for abundance of Type IIa/IIx-2 myonuclei, the presence of which appeared to be almost entirely dependent on MuSCs, suggesting these are myonuclei that had fused in during MOV. (Figure 1J-M).

### Differential responses to MOV in adult mice with and without MuSCs revealed the gene signature of MuSC-Derived myonuclei and MOV-Responsive resident myonuclei

A major strength of being able to deplete MuSCs is it enables identification of MuSC-Derived myonuclei in response to MOV. Using Density Based Spatial Clustering of Applications with Noise (DBSCAN)^9^ which allows delineation of myonuclear clusters based on differential density, we identified a high-density cluster enriched with myonuclei from Adult MOV MuSC**^+^** (Log2 Fold Enrichment = 2.12, 21.8% of myonuclei in Adult MOV MuSC**^+^**) which we defined as MuSC-Derived myonuclei (Figure 2A-C). This cluster also appeared to overlap with the Type IIa/IIx cluster identified in Figure 1J-M. Plotting the density map across 3 dimensions (UMAP1, UMAP 2, differential density) revealed the distinct shape of this region (Figure S1A). We used Lasso, a regression model optimized for sparse data feature selection^10^ to determine genes most important for shaping differential density. Lasso identified a 323 gene model to reconstruct the differential density map between Adult MOV MuSC**^+^** and Adult MOV MuSC**^-^** (r = 0.937, R^2^ = 0.835; Figure S1A-B) which defined the gene signature of MuSC-Derived myonuclei. From the 323 gene model, 161 genes had positive Lasso coefficients which represent genes that best predict high density and are most specific to MuSC-Derived myonuclei. The top 50 genes by Lasso coefficient include the lncRNA, *H19*, a previously identified marker of MuSC-Derived myonuclei^11^ and a transcription factor, *Bnc2* (Figure 1D). In addition, ENCODE/ChEA transcription factor target enrichment revealed transcription factors targeting genes most specific to MuSC-Derived myonuclei (positive lasso coefficients) including *Myod1*, *Smc3*, *E2f1*, *Creb1*, *Tcf3*, *Ubtf*, *Pbx3*, and the Yamanaka factor, *Myc* which is induced with hypertrophic stimuli^12–14^ (Figure 2E). Similar to the top genes identified using Lasso, “traditional” differential expression-based marker genes for this cluster also included *H19* and *Bnc2* alongside *Runx1*, *Ncam1*, and *Map1b* (Figure 2F). Gene Ontology Biological Process (GOBP) enrichment for these marker genes included Muscle Contraction, Regulation of Supramolecular Fiber Organization, Positive Regulation of DNA-Templated Transcription, Intracellular Signaling Cassette, and Positive Regulation of Axonogenesis (Figure 2G).

**Figure 2:**
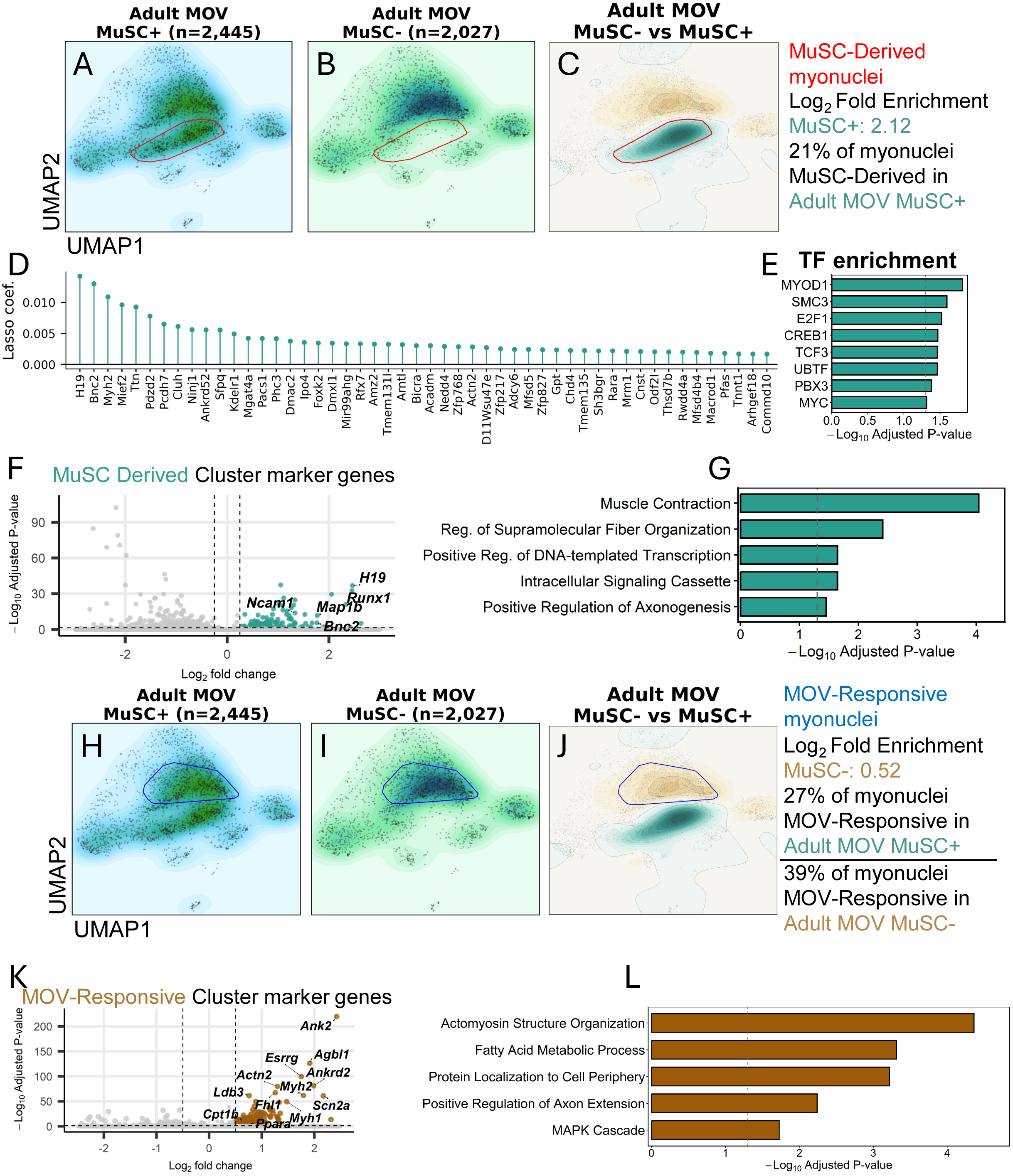
MuSC derived and MOV-Responsive myonuclei are distinct myonuclear populations. A. Count-normalized density of myonuclear abundance within the integrated UMAP from Adult MOV MuSC**^+^** and B. Adult MOV MuSC**^-^** samples with convex hull indicating the satellite cell derived myonuclear cluster. C. Differential normalized density between Adult MOV MuSC**^+^** and Adult MOV MuSC**^-^** samples with convex hull indicating the satellite cell derived myonuclear cluster. D. Top 50 high density (positive Lasso coefficients) predicting genes. E. Transcription factor enrichment of gene targets from *D*. F. Differential expression of genes within the positive differential density (Adult MOV MuSC**^+^** biased) convex hull compared to all other myonuclei. G. Enriched Gene Ontology Biological Processes of positively differentially expressed genes (marker genes) from *F.* H. Count-normalized density of myonuclear abundance within the integrated UMAP from Adult MOV MuSC**^+^**and I. Adult MOV MuSC**^-^**samples with convex hull indicating the MOV-Responsive resident myonuclei cluster. J. Differential normalized density between Adult MOV MuSC**^+^** and Adult MOV MuSC**^-^** samples with convex hull indicating the MOV-Responsive resident myonuclei cluster. K. Differential expression of genes within the negative (Adult MOV MuSC**^-^** biased) differential density convex hull compared to all other myonuclei. L. Enriched Gene Ontology Biological Processes of differentially expressed genes from *K*.

Having characterized MuSC-Derived myonuclei, we next focused on characterizing the response of resident (i.e. non-satellite cell-derived) myonuclei to MOV in Adult muscle with or without MuSCs. These MOV-Responsive myonuclei resided in a distinct region enriched with myonuclei from both Adult MOV MuSC**^+^** and Adult MOV MuSC**^-^** mice (MuSC**^-^** Log_2_ Fold Enrichment = 0.52, 27% of myonuclei in Adult MOV MuSC**^+^**, 39% of myonuclei in Adult MOV MuSC**^-^**; Figure 2H-J). Greater relative abundance of MOV-Responsive resident myonuclei in MuSC-depleted muscle may reflect a compensatory response to MOV without MuSCs. This is consistent with previous findings showing myonuclei have an ability to increase transcriptional output without an increase in myonuclear abundance^15^ (PMID: 26764089). Unlike identification of MuSC-Derived myonuclei, differential density within this region was low and more uniformly distributed (Figure S1E). As a result, Lasso regression failed to define a gene set that accurately reconstructed differential density biased toward Adult MOV MuSC**^-^** (Figure S1F). Resident myonuclear transcriptional responses to MOV are therefore not meaningfully affected by MuSC-depletion in Adult muscle. These myonuclei were designated as “MOV-Responsive” resident myonuclei (Figure 2H-J). Using marker gene expression of MOV-Responsive resident myonuclei to uncover their defining gene signature, we found *Ank2*, *Agbl1*, *Ankrd2*, *Ldb3*, and *Fhl1* that are critical for supporting muscle structural integrity, alongside *Esrrg*, *Cpt1b*, and *Ppara* that are important for aerobic and fatty acid metabolic adaptation (Figure 2K). GOBP enrichment revealed this cluster is also enriched in processes including Actomyosin Structure Organization, Fatty Acid Metabolic Process, Protein Localization to Cell Periphery, Positive Regulation of Axon Extension and MAPK Cascade (Figures 2L).

### Aged myonuclei have a diminished response to MOV compared to Adult myonuclei that is exacerbated with MuSC depletion

Next, we determined the effect of MuSC depletion in Aged muscle by comparing Aged MOV MuSC**^+^** to Aged MOV MuSC**^-^**conditions. We performed differential density between Aged MOV MuSC**^+^** (Figure 3A) and Aged MOV MuSC**^-^** myonuclei (Figure 3B). We identified two distinct high-density clusters enriched with myonuclei from Aged MOV MuSC**^+^** muscle, similar to the resident myonuclear MOV-Responsive gene signatures (MuSC**^+^** Log_2_ Fold Enrichment = 0.66, 24.85% of myonuclei in Aged MOV MuSC**^+^**) and MuSC-Derived (MuSC**^+^** Log_2_ Fold Enrichment = 2.74, 8.09% of myonuclei in Aged MOV MuSC**^+^**; Figure 3A-C). Two clusters that were enriched in Aged MOV MuSC**^-^** myonuclei over Aged MOV MuSC**^+^** myonuclei (Cluster 1 Log_2_ Fold Enrichment = 0.92; Cluster 2 Log_2_ Fold Enrichment = 0.72; Figure 3D) overlapped with regions enriched with myonuclei from MuSC**^+^** and MuSC**^-^** Sham muscle (26% within MOV-enriched region, 74% within Sham-enriched region; Figure 3E-F). The defining feature of MuSC presence in Aged MOV muscle was greater abundance of both MOV responsive myonuclei and MuSC derived myonuclei. This suggests transcriptional contributions from these myonuclear populations are dependent on MuSCs in Aged muscle. Further, Aged MOV MuSC**^-^**muscle was enriched with myonuclei that overlapped with Sham muscle, suggesting Aged MOV MuSC**^-^**myonuclei mount a diminished transcriptional response to MOV compared to Aged MOV MuSC**^+^**myonuclei.

**Figure 3:**
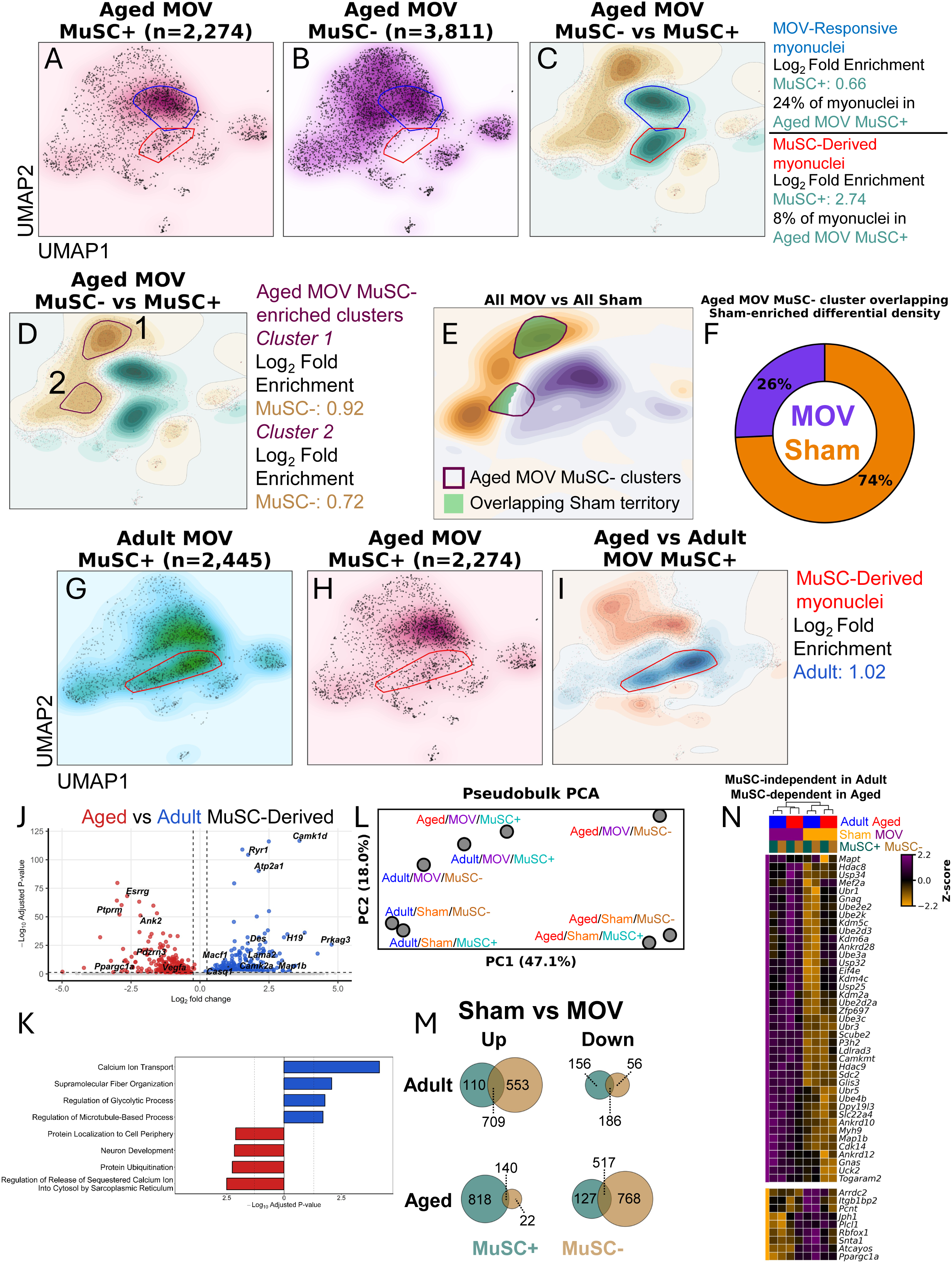
SC depletion impairs transcriptional responsiveness to MOV in Aged mice alongside lower abundance of MuSC Derived myonuclei with MOV compared to Adult mice. A. Count-normalized density of myonuclear abundance within the integrated UMAP from Aged MOV MuSC**^+^**and B. Aged MOV MuSC**^-^**samples. C. Differential density between Aged MOV MuSC**^+^** and Aged MOV MuSC**^-^** with convex hulls indicating previously identified MuSC-Derived myonuclei and MOV-Responsive resident myonuclei superimposed with convex hulls identifying clusters of differential density biased toward Aged MOV MuSC**^+^** myonuclei compared to Aged MOV MuSC**^-^**. D. Two distinct convex hulls identifying Aged MOV MuSC**^-^** enriched differential density regions. E. Differential density comparing All MOV and Sham myonuclei with convex hulls identified in *D* and filled where they overlap with Sham-enriched regions. F. Distribution of Aged MOV MuSC**^-^**myonuclei overlapping Sham-enriched territories within convex hulls identified in *D*. G. count-normalized density maps for Adult MOV MuSC**^+^** and H. Aged MOV MuSC**^+^** with convex hulls for MuSC-Derived myonuclei and I. differential density enriched with nuclei from Adult MOV MuSC**^+^**. J. Differential expression of Adult vs Aged MuSC-Derived myonuclei. K. Biological Process pathway enrichment of differentially expressed genes in *J*. L. Principal component analysis of individual samples in pseudobulk. M. Venn diagrams showing number of genes up-and down-regulated in Sham vs MOV comparisons for Adult MuSC+, Adult MuSC-, Aged MuSC+, and Aged MuSC-samples.

### MuSC-Derived myonuclei constitute the greatest age-dependent myonuclear transcriptional response to MOV

Differential density between Aged MOV MuSC**^+^** and Adult MOV MuSC**^+^** myonuclei revealed enrichment of MuSC-Derived myonuclei in Adult MOV MuSC**^+^**over Aged MOV MuSC**^+^**(Adult Log_2_ Fold Enrichment = 1.02). MuSC-Derived myonuclei are therefore more abundant in Adult MOV MuSC**^+^** muscle than in Aged MOV MuSC**^+^** muscle (Figure 3G-I). Differential expression between Adult and Aged myonuclei within this cluster revealed enrichment of GOBPs Calcium Ion Transport, Supramolecular Fiber Organization, Regulation of Glycolytic Process and Regulation of Microtubule-Based Process in Adult myonuclei and enrichment of Regulation of Release of Sequestered Calcium Ion Into Cytosol by Sarcoplasmic Reticulum, Protein Ubiquitination, Neuron Development, and Protein Localization to Cell Periphery in Aged muscle (Figure 3J-K), suggesting differential functions of MuSC-Derived myonuclei in Adult and Aged muscle. Principle component analysis of pseudobulked snRNA-seq data revealed PC1, presumably age, accounted for 47% of the variability while PC2, presumably MOV, accounted for 18% of the variability. Consistent with our previous studies^6,16^, regardless of age, the depletion of MuSCs did not have much impact under resting conditions, i.e., Sham (Figure 3L). The most striking finding from the PC analysis was that MOV dramatically shifted the Aged group transcriptionally to be more similar, i.e., closer, to the Adult group only when MuSCs were present (Figure 3L), thus supporting MuSC-dependent transcriptional rejuvenation of Aged myonuclei^17^. In addition, differentially expressed genes in response to MOV (Sham vs MOV) were only modestly different between Adult MuSC**^+^** and Adult MuSC**^-^** myonuclei (Figure 3M). In Aged muscle however, Aged MuSC**^+^**had far more uniquely upregulated genes than Aged MuSC**^-^** in response to MOV while Aged MuSC**^-^** had far more uniquely downregulated genes (Figure 3M). Genes selected by MuSC-independent differential expression with MOV in Adult myonuclei (shared between Adult MOV MuSC**^+^** vs Adult Sham MuSC**^+^** and Adult MOV MuSC**^-^** vs Adult Sham MuSC**^-^**) and MuSC-dependent differential expression in Aged muscle (differentially expressed only in Aged MOV MuSC**^+^** vs Aged Sham MuSC**^+^**) included upregulation of microtubule-related genes (*Map1b*, *Mapt*), Ankrd family genes (*Ankrd10*, *Ankrd12*, *Ankrd28*), G-protein coupled receptor signaling genes (*Gnaq*, *Gnas*), histone deacetylases (*Hdac8*, *Hdac9*), lysine demethylases (*Kdm2a*, *Kdm4c*, *Kdm5c*, *Kdm6a*), and ubiquitin-based protein quality control (*Ube2d2a*, *Ube2d3*, *Ube2e2*, *Ube2k*, *Ube3a*, *Ube3c*, *Ube4b*, *Ubr1*, *Ubr3*, *Ubr5*, *Usp25*, *Usp32*, *Usp34*) and notably, downregulation of the master regulator of mitochondrial biogenesis, *Ppargc1a* (Figure 3N). This suggests the role of MuSCs in Aged muscle adaptation to MOV is generally targeted toward microtubule remodeling, stress response via *Ankrd* family gene expression, G-protein coupled receptor signaling, chromatin remodeling, and protein quality control. By contrast, these responses are not dependent on MuSCs in Adult muscle.

### MuSC-Derived myonuclei preferentially differentiate toward NMJ, MTJ, and Spindle myonuclear signatures in response to MOV

To define the fate of MuSC-Derived myonuclei, Diffusion Pseudotime transcriptional trajectory analysis was performed on the full MuSC**^+^**subset of myonuclei (Adult and Aged, MOV and Sham, MuSC**^+^** and MuSC**^-^** samples) with the root node defined as the centroid of the MuSC-Derived myonuclear cluster (Figure 4A-C). The highest (most differentiated) pseudotime values were assigned to myonuclei belonging to myotendinous junction (MTJ), neuromuscular junction (NMJ), and Spindle (intrafusal fiber) myonuclear clusters (Figure 4B-C). Using CellRank’s pseudotime kernel to calculate terminal states^18,19^, we confirmed that terminal transcriptional states represented Spindle, MTJ, and NMJ myonuclear clusters (Figure 4D). To identify myonuclei undergoing the greatest rate of transcriptional shift, a fate commitment gradient was calculated using per-nucleus terminal state bias (absorption probabilities toward each terminal state). The fate commitment gradient identified the most rapid transcriptional shift from MuSC-Derived myonuclei toward Spindle, MTJ, and NMJ clusters (Figure 4E-F), predicting MuSC-Derived myonuclei undergo rapid transcriptional shifts toward these highly specialized myonuclear clusters.

**Figure 4.**
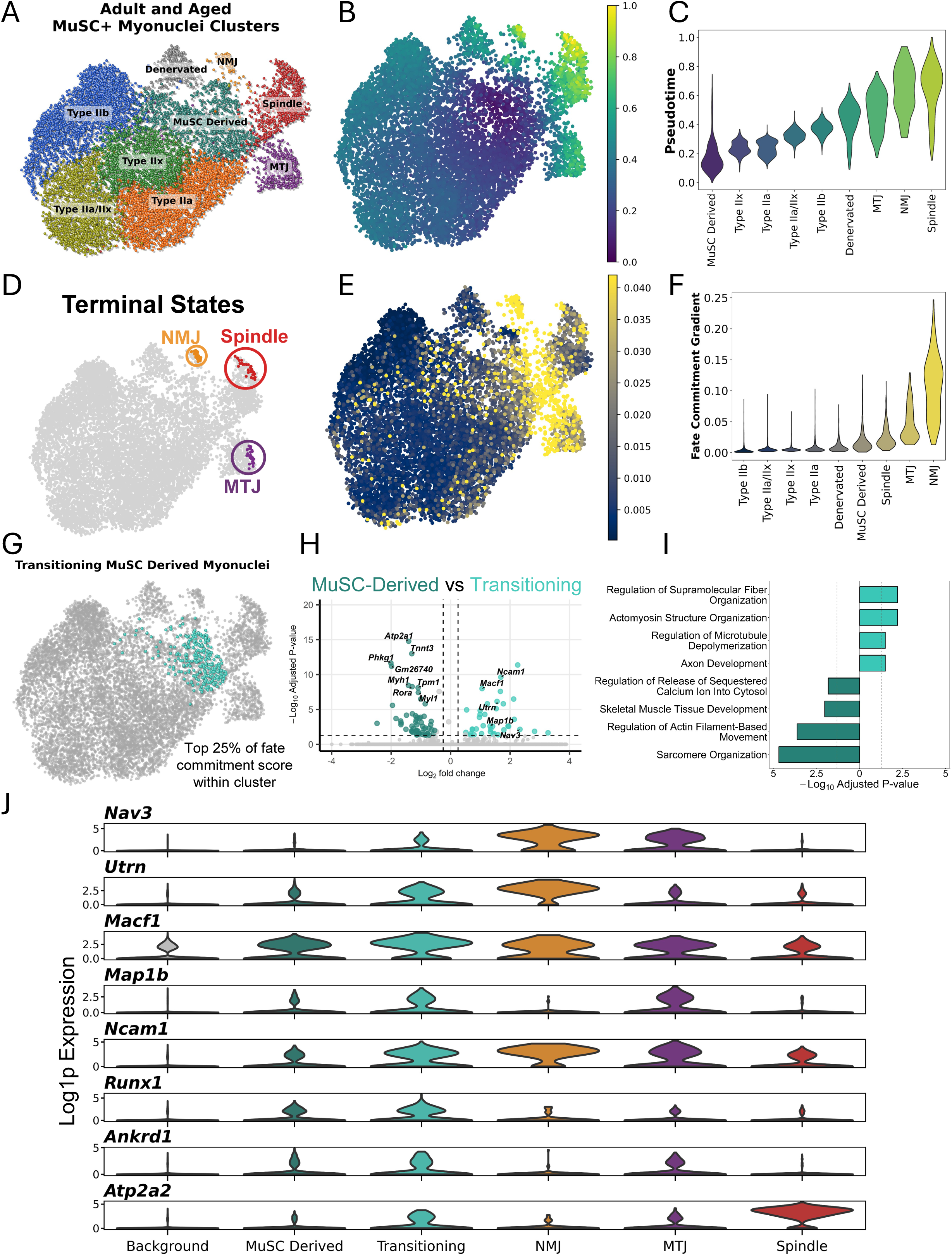
MuSC Derived myonuclei transcriptionally specialize toward Spindle, MTJ, and NMJ myonuclear transcriptional signatures. A. UMAP plot of myonuclei from MuSC**^+^** muscle only. B. Diffusion pseudotime trajectory with the centroid of the MuSC-Derived myonuclear cluster as the root node. C. Distribution of pseudotime across myonuclear subtypes. D. Terminal states of MuSC-Derived myonuclei E. Fate commitment gradient of myonuclei identifying myonuclei most likely to transcriptionally shift toward a specific terminal state. F. Distribution of fate commitment gradient scores across myonuclear clusters. G. Transitioning MuSC Derived myonuclei identified as nuclei within the top 25% of fate commitment gradient scores. H. Differential expression between Transitioning myonuclei identified in *G* and MuSC-Derived myonuclei. I. Biological Process pathway enrichment from differentially expressed genes in *H*. J. Gene expression pattern of genes significantly correlating with Fate Commitment Gradient.

To understand the transcriptional signature of MuSC-Derived myonuclei responsible for this transcriptional shift, we defined “Transitioning MuSC-Derived myonuclei” as the top 25% of the fate commitment gradient scores within the MuSC-Derived myonuclear cluster, representing MuSC-Derived myonuclei most likely to change transcriptional states (Figure 4G). Differential expression in these myonuclei compared to all other MuSC-Derived myonuclei (Figure 4H) revealed enriched GOBPs including Regulation of Supramolecular Fiber Organization, Actomyosin Structure Organization, Regulation of Microtubule Depolymerization, and Axon Development alongside MuSC-Derived myonuclei being enriched for Sarcomere Organization, Regulation of Actin Filament-Based Movement, Skeletal Muscle Tissue Development, and Regulation of Release of Sequestered Calcium Ion Into Cytosol (Figure 4I). Top differentially expressed genes enriched in Transitioning MuSC-Derived myonuclei compared to all other MuSC-Derived myonuclei included the microtubule plus-end tracking (+TIP) gene, *Nav3* and microtubule remodeling genes, *Macf1,* which is known to facilitate physical myonuclear translocation^20^, and *Map1b* (Figure 4H, 4J). In Spindle myonuclei, the slow muscle SERCA, *Atp2a2* increased in expression in Transitioning myonuclei and is a very specific and highly expressed gene in Spindle myonuclei. In addition, *Runx1* was more highly expressed in the transitioning myonuclei compared to other MuSC-Derived myonuclei (Figure 4J), a cluster for which *Runx1* was already a defining gene (Figures 2F, 2M). These genes also increased in expression among MuSC-Derived myonuclei as they progressed from MuSC-Derived to Transitioning myonuclear clusters. *Macf1* and *Ncam1* specifically were expressed not only in the MuSC-Derived and Transitioning clusters, but also in each terminal state cluster (NMJ, MTJ, and Spindle clusters). *Nav3* expression was specific to NMJ and MTJ terminal states. *Utrn* expression was specific to NMJ while *Map1b* and the myonuclear stress-response gene, *Ankrd1* were specific to MTJ terminal state, and *Atp2a2* was specific to the Spindle cluster (Figure 4J). Interestingly, although *Runx1* expression increased from MuSC-Derived to Transitioning myonuclei, *Runx1* expression was restricted to MuSC-Derived and Transitioning myonuclei (Figure 4J), suggesting it may regulate a general post-fusion myonuclear specialization program independent of a terminal state-specific target.

### Aging divergently delineates MOV-Responsive MTJ and NMJ transcriptional specialization of newly fused MuSC-Derived myonuclei

To determine transcriptional fate progression of Adult MuSC-Derived myonuclei in response to MOV, myonuclei from Adult MOV MuSC**^+^**and Adult Sham MuSC**^+^**were subsetted for Diffusion Pseudotime and subsequent Fate Commitment analyses with the root node identified as the centroid of the MuSC-Derived myonuclear cluster (Figure 5A-E). The highest calculated pseudotime values were assigned to myonuclei belonging in Spindle and MTJ myonuclear clusters (Figure 5B-C). Similarly, the highest fate commitment scores were also assigned to myonuclei belonging to Spindle and MTJ clusters (Figure 5D-E). In addition, terminal state analysis revealed transcriptional endpoints of MuSC-Derived myonuclei in MTJ and Spindle myonuclear clusters (Figure 5F). To determine the effect of aging, we performed a parallel analysis on the same subset of myonuclei from Aged muscle (Aged MOV MuSC**^+^**, Aged MOV MuSC**^-^**, Figure 5G-L). Similar to Adult muscle, MuSC-Derived myonuclei in Aged muscle had a pseudotime transcriptional trajectory toward Spindle myonuclei in response to MOV. However, unlike Adult muscle, the highest pseudotime values were assigned to NMJ myonuclei (Figure 5H-I). In addition, the highest fate commitment scores in Aged muscle were also assigned to myonuclei belonging to the NMJ cluster (Figure 5J-K). Although Aged MuSC-Derived myonuclei had terminal states in NMJ and Spindle myonuclear clusters, an additional terminal transcriptional state within the MuSC-Derived myonuclear cluster was also identified in Aged muscle only (Figure 5L), suggesting a potentially impaired ability of Aged MuSC-Derived myonuclei to transcriptionally specialize post-fusion.

**Figure 5:**
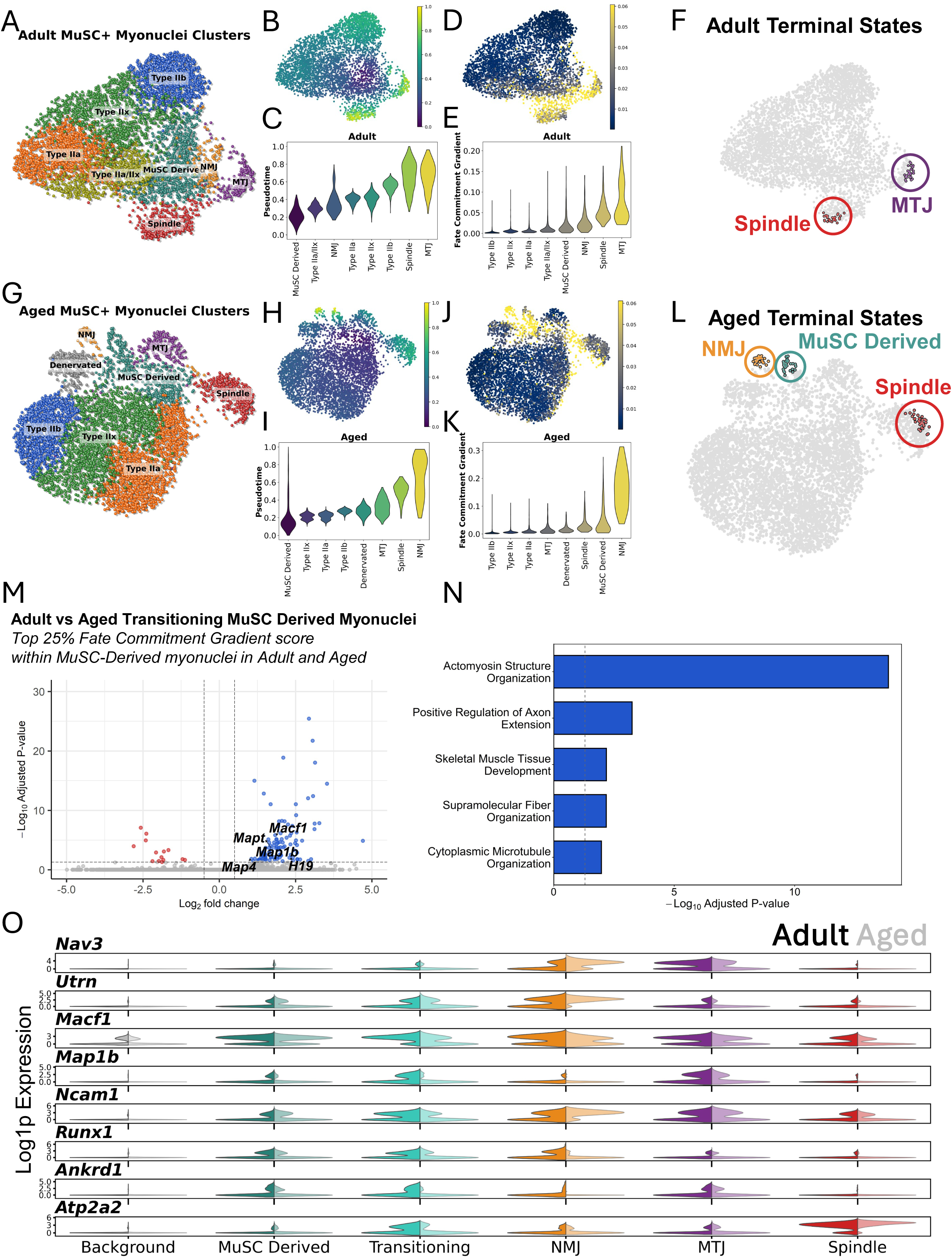
Adult and Aged MuSC Derived myonuclei specialize toward MTJ and NMJ transcriptional programs. A. UMAP of Adult MOV MuSC**^+^**and Adult Sham MuSC**^+^** (Adult MuSC**^+^**) myonuclei. B. Diffusion pseudotime trajectory of Adult MuSC**^+^** myonuclei. C. Distribution of pseudotime across Adult MuSC**^+^**myonuclear subtypes. D. Fate commitment gradient of Adult MuSC**^+^**myonuclei scoring likelihood of myonuclei transcriptionally shifting toward a specific terminal state. E. Distribution of fate commitment gradient scores across Adult MuSC**^+^** myonuclear clusters. F. Terminal state prediction of Adult MuSC**^+^** MuSC Derived myonuclei. G. UMAP of Aged MOV MuSC**^+^** and Aged Sham MuSC**^+^** (Aged MuSC**^+^**) myonuclei. H. Diffusion pseudotime trajectory of Aged MuSC**^+^** myonuclei. I. Distribution of pseudotime across Aged MuSC**^+^**myonuclear subtypes. J. Fate commitment gradient of Aged MuSC**^+^** myonuclei scoring likelihood of myonuclei transcriptionally shifting toward a specific terminal state. K. Distribution of fate commitment gradient scores across Aged MuSC**^+^** myonuclear clusters. L. Terminal state prediction of Aged MuSC**^+^** MuSC Derived myonuclei. M. Differential expression between Adult and Aged transitioning MuSC Derived myonuclei defined as the top 25% of fate commitment gradient scores in Adult MuSC**^+^** and Aged MuSC**^+^** myonuclei. N. Biological Process pathway enrichment and differential expression of genes from *M*. O. Gene expression pattern of genes significantly correlating with Fate Commitment Gradient.

To determine the effect of advanced age on the myonuclear transcriptional fate of MuSC-Derived myonuclei, the top 25% of fate commitment scores within MuSC-Derived myonuclei for Adult and Aged myonuclei were used to define Transitioning myonuclei similar to those defined in Figure 4G. Differential expression between Adult and Aged Transitioning myonuclei revealed most differentially expressed genes were enriched in Adult over Aged Transitioning myonuclei and included genes that are associated with microtubule remodeling: *Macf1*, *Map1b*, *Mapt*, and *Map4* (Figure 5M). Differential expression also revealed GOBP enrichment in Adult Transitioning myonuclei including Actomyosin Structure Organization, Positive Regulation of Axon Extension, Skeletal Muscle Tissue Development, Supramolecular Fiber Organization, and Cytoplasmic Microtubule Organization, while no GOBPs were enriched in Aged Transitioning myonuclei (Figure 5N). *Nav3* showed preferential expression in MTJ myonuclei in Adult muscle, while in Aged muscle, *Nav3* expression was biased toward NMJ myonuclei (Figure 5O) suggesting it may play a role in guiding terminal transcriptional specialization of MuSC-Derived myonuclei. *Nav3* is known to guide microtubule polymerization and has a well described and highly conserved role in promoting synaptic formation in the central nervous system^21–23^, and is expressed in human MTJ^24^. *Runx1* expression was higher in Adult than in Aged MuSC-Derived and Transitioning myonuclei and this observation was accompanied by higher expression of *Macf1*, *Map1b*, and *Ncam1* in Adult compared to Aged myonuclei (Figure 5O).

### Runx1 represses MuSC-Derived myonuclear differentiation and Esrrg transcriptionally specializes MuSC-Derived myonuclei in intrafusal Spindle fibers

We applied CellOracle^25^, which predicts the effect of *in-silico* transcription factor (TF) knockout (KO) on differentiation trajectories inferred by pseudotime. CellOracle assigns inner products to UMAP regions that score whether knockout of a specific transcription factor promotes differentiation (positive inner product) or represses differentiation (negative inner product). Relating to pseudotime, positive inner products assigned to a UMAP region indicates the transcription factor in question represses differentiation to the transcriptional identity in that region while negative inner products indicate the transcription factor promotes differentiation to that region.

To determine gene drivers of transcriptional differentiation of MuSC-Derived myonuclei toward specific transcriptional states, MuSC-Derived myonuclei were subclustered and assigned a trajectory target based on their absorption probability bias computed by CellRank. In Adult muscle, MTJ-targeted and Spindle-targeted MuSC-Derived myonuclei represented distinct myonuclear populations in transcriptional UMAP space (Figure 6A). MTJ-targeted myonuclei were marked by higher expression of *Cbfb,* RUNX1’s obligate DNA binding cofactor, while Spindle-targeted myonuclei were marked by higher expression of *Esrrg* (Figure 6B). To determine predicted roles of *Runx1*, *Cbfb*, and *Esrrg* in MuSC-Derived differentiation trajectories, we found *Runx1*, *Cbfb*, and *Esrrg* all played generally repressive roles in myonuclear transcriptional differentiation (Figure 6C). When restricting differentiation trajectory to the MTJ cluster, *Runx1* and *Cbfb* were predicted to be permissive for MuSC-Derived myonuclear differentiation (Figure 6D-E). Conversely, *Runx1* and *Cbfb* were restrictive against differentiation toward other clusters (Figure 6D-E). In addition, *Runx1* expression in Adult was almost exclusively restricted to the MuSC-Derived myonuclear cluster, suggesting a potential role of RUNX1 in MuSC-Derived myonuclear specialization (Figure 6F). Similarly, *Esrrg* was permissive for differentiation of MuSC-Derived myonuclei toward Spindle, but restrictive against differentiation to other clusters (Figure 6G-H). In addition, both *Runx1* and *Cbfb* were predicted to repress Spindle differentiation of MuSC-Derived myonuclei, suggesting Runx1 may delineate MTJ over Spindle differentiation of MuSC-Derived myonuclei in Adult muscle (Figure 6G). Unlike *Runx1*, *Esrrg* expression was more ubiquitous across more oxidative clusters (Type IIa, Type IIa/IIx) alongside MuSC-Derived and Spindle myonuclei, consistent with previous observations of *Esrrg* expression in slow/oxidative muscle and muscle fibers^26,27^ (Figure 6I).

**Figure 6:**
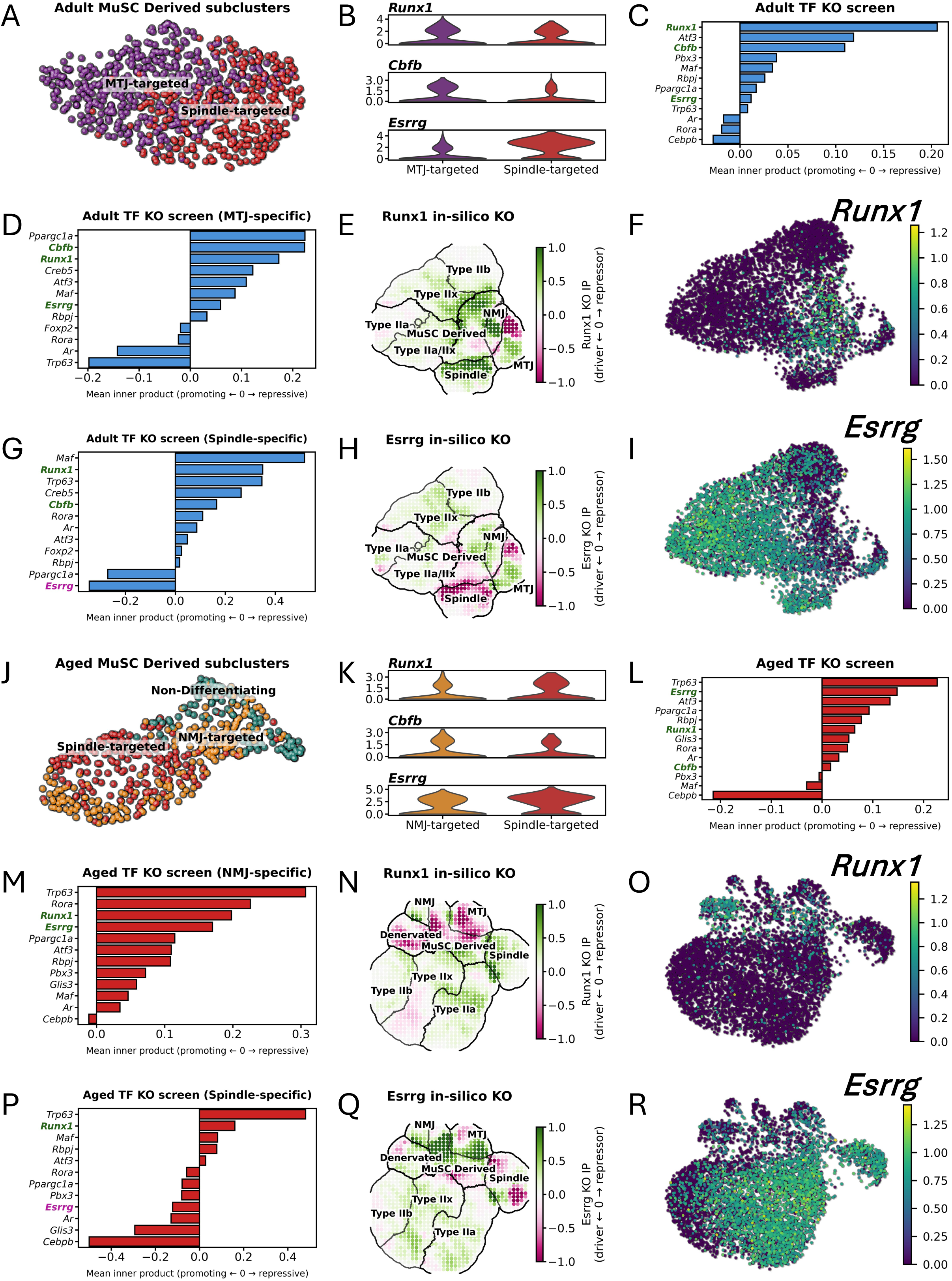
Runx1 represses differentiation of MuSC-Derived myonuclei while Esrrg promotes Spindle-specific differentiation of MuSC-Derived myonuclei. A. UMAP plot of MTJ-targeted and Spindle-targeted MuSC-Derived myonuclei in Adult muscle. B. Expression profiles of *Runx1*, *Cbfb*, and *Esrrg* in MTJ-targeted and Spindle-targeted MuSC-Derived myonuclei in Adult muscle. C. In-silico transcription factor (TF) KO screen, ranked by mean inner product across the full UMAP space; negative values are assigned TFs that promote the pseudotime trajectory while positive values are assigned to TFs that repress the pseudotime trajectory. D. In-silico TF KO screen with inner products for TFs across MTJ-targeted and MTJ myonuclei in Adult muscle. E. Grid map of inner products for predicted Runx1 KO across the Adult MuSC+ (Adult MOV MuSC+ and Adult Sham MuSC+) UMAP. F. Feature plot of *Runx1* expression in Adult myonuclei. G. In-silico TF KO screen with inner products for TFs across Adult Spindle-targeted and Spindle myonuclei. H. Grid map of inner products for predicted Esrrg KO across the Adult MuSC+ (Adult MOV MuSC+ and Adult Sham MuSC+) UMAP. I. Feature plot of *Esrrg* expression in Adult myonuclei. J. UMAP plot of Spindle-targeted, NMJ-targeted, and Non-differentiating MuSC-Derived myonuclei in Aged muscle. K. Expression profiles of *Runx1*, *Cbfb*, and *Esrrg* in NMJ-targeted and Spindle-targeted MuSC-Derived myonuclei in Aged muscle. L. In-silico transcription factor (TF) KO screen, ranked by mean inner product across the full UMAP space. M. In-silico TF KO screen with inner products for TFs across NMJ-targeted and NMJ myonuclei in Aged muscle. N. Grid map of inner products for predicted Runx1 KO across the Aged MuSC+ (Aged MOV MuSC+ and Aged Sham MuSC+) UMAP. O. Feature plot of *Runx1* expression in Aged myonuclei. P. In-silico TF KO screen with inner products for TFs across Aged Spindle-targeted and Spindle myonuclei. Q. Grid map of inner products for predicted Esrrg KO across the Aged MuSC+ (Aged MOV MuSC+ and Aged Sham MuSC+) UMAP. R. Feature plot of *Esrrg* expression in Aged myonuclei.

In Aged muscle, Spindle and NMJ were identified as terminal transcriptional states of MuSC-Derived myonuclei, and similar to Adult, MuSC-Derived myonuclei were assigned as targeting their predicted terminal transcriptional states (Figure 6J). *Runx1* expression is modestly higher in Spindle-targeted myonuclei while *Cbfb* is higher in NMJ-targeted myonuclei and consistent with Adult, *Esrrg* is higher in Spindle-targeted than NMJ-targeted myonuclei (Figure 6K). Similar to Adult, *Esrrg*, *Runx1*, and *Cbfb* were all predicted to generally repress differentiation of MuSC-Derived myonuclei (Figure 6L). In Aged muscle, *Runx1* was also the top predicted transcription factor to inhibit differentiation of MuSC-Derived myonuclei toward the NMJ cluster while promoting differentiation toward Denervated and MTJ clusters (Figure 6M-N). Similar to Adult, *Runx1* expression was high in MuSC-Derived myonuclei and in Aged only, *Runx1* was also expressed in Denervated myonuclei (Figure 6O). When restricting inner products to the Spindle trajectory, *Esrrg* is also a driving transcription factor of the Spindle trajectory, albeit a weaker driver of Spindle differentiation than in Adult myonuclei (Figure 6P-Q). Similar to Adult, *Esrrg* was widely expressed across multiple myonuclear types (Figure 6R), but in-silico KO of *Esrrg* revealed its role as a driving transcription factor for MuSC-Derived myonuclear differentiation toward Spindle myonuclei.

These in-silico predictions establish evidence of *Runx1’s* role as generally repressing differentiation of MuSC-Derived myonuclei while *Esrrg* is a transcriptional driver of specialization for the rare population of MuSC-Derived myonuclei that exist only within intrafusal fibers.

## Discussion

Using snRNA-seq of enriched myonuclei in young Adult and Aged mechanically overloaded skeletal muscle with and without MuSCs we delineated terminal transcriptional states of MuSC-Derived myonuclei as MTJ, NMJ, and Spindle myonuclear identities in Adult and Aged muscle, respectively. We also found that the transcriptional response of Aged muscle to MOV is diminished compared to that of young adult muscle and that MuSCs are necessary for myonuclear transcriptional responsiveness to MOV in Aged muscle.

Using differential density analysis combined with the use of Lasso regression-based feature selection, we defined the gene signature of MuSC-Derived myonuclei. Our analytical approach was biologically confirmed by *H19* being the top gene defining the highest density region of MuSC-Derived myonuclei, in agreement with the Millay lab^11^. Using the comparison between MOV Adult myonuclei from muscle with or without MuSCs, we identified the transcriptional signature of resident myonuclei that are responsive to MOV. In Adult muscle, these myonuclei were not dramatically different in biased enrichment toward muscle with or without MuSCs, with appreciable abundance from both groups. This myonuclear population was also distinctly different from MuSC-Derived myonuclei, as enriched expression of genes associated with metabolic remodeling in response to MOV was apparent, while the MuSC-Derived myonuclei were enriched with microtubule-associated genes alongside *Runx1* and *H19*. These data suggest that in Aged muscle, MuSC presence is important for both MuSC-Derived myonuclear abundance and compensation of impaired resident myonuclear responsiveness to MOV. In Adult muscle, MuSC presence appears to only contribute to MuSC-Derived myonuclear abundance, while the resident myonuclei are unaffected by MuSC-depletion suggesting MuSC fusion does not influence resident myonuclear responsiveness at 14-days of MOV, studied here. With a more prolonged hypertrophic stimulus however, MuSCs appear to become important, even in Adult muscle^28,29^.

In Aged myonuclei, we found the transcriptional responsiveness to MOV is diminished, consistent with previous studies showing impaired response of aged muscle to resistance exercise^2,3^. We showed the pseudobulk transcriptional profile of myonuclei from overloaded Aged muscle with MuSCs as being dramatically different from all other Aged muscle samples such that they are “closer” to Adult overloaded muscle and thereby transcriptionally “younger” in transcriptional PCA space. The MuSC-and MOV-dependent transcriptional proximity of Aged myonuclei to Adult myonuclei represents a MuSC activity-dependent reversal of the age-associated myonuclear transcriptional profile. Differential density between Aged mice with and without MuSCs during MOV revealed MuSC-dependent myonuclear transcriptional responsiveness to MOV in advanced age. We show that in advanced age, myonuclei of MuSC-depleted muscle are biased toward transcriptional space that is not aligned with either resident MOV-Responsive or MuSC-Derived myonuclear clusters. Conversely, enrichment of myonuclei in both resident MOV-Responsive and MuSC-Derived clusters depended on the presence of MuSCs in Aged muscle. This suggests MuSC-Derived myonuclei can compensate for lost responsiveness of resident myonuclei in Aged muscle. In addition, differential density between Adult and Aged myonuclei during MOV revealed the greatest transcriptional difference between the two ages is the presence of MuSC-Derived myonuclei. This demonstrates MuSC-Derived myonuclei as the central node of the youthful myonuclear transcriptional signature.

A major finding from our snRNA-seq analysis was the transcriptional fate of MuSC-Derived myonuclei. These myonuclei specialized toward the MTJ, NMJ, and Spindle fiber myonuclear transcriptional signatures in overloaded Adult and Aged muscle, respectively. Using trajectory inference analysis tools with MuSC-Derived myonuclei as the root node of the trajectory, we identified a subset of MuSC-Derived myonuclei as actively transitioning toward specialized myonuclear transcriptional signatures. These transitioning myonuclei were enriched with genes associated with microtubules and microtubule remodeling. Specifically, *Macf1* was strikingly enriched in these myonuclei and is known to govern myonuclear translocation to support NMJ integrity^20^. Other cluster-defining genes known to associate with microtubule remodeling included *Map1b*, which specifically associates with acetylcholine receptors at the NMJ^20^ but our data suggests *Map1b* expression is MTJ-specific. MuSC-Derived and Transitioning myonuclei were also enriched with the transcription factor *Runx1*, which is broadly known to associate with a denervation response, but the exact function or target genes of *Runx1* in skeletal muscle remain unknown. Recently, *Runx1* has been identified as part of the hypertrophic response^8,30^ and regulates muscle mass, myofibrillar organization and autophagy in skeletal muscle^31^. Given these prior observations and our data here, we posit *Runx1* may be a critical regulator of the transition state of MuSC-Derived myonuclei. Advanced age appeared to affect the trajectory of these transitioning myonuclei, where young Adult transitioning myonuclei had a differentiation trajectory ending with an MTJ-specialized transcriptional signature, while Aged transitioning myonuclei specialized toward an NMJ myonuclear transcriptional signature and a Spindle-directed trajectory in both ages. The transcriptional trajectory of the transitioning myonuclei included a terminal state within the MuSC-Derived cluster only in Aged muscle, which we interpret as age-induced impairment of post-fusion myonuclear transcriptional specialization. We also observed in aged myonuclei lower expression of the microtubule genes that defined the transitioning myonuclei, including *Macf1*, *Map1b*, *Mapt*, and *Map4*. Lower expression of these microtubule genes within the transitioning MuSC-Derived myonuclei may underlie previous reports of impaired microtubule dynamics^32,33^ that contribute to age-associated resistance to adaptation and delayed transcriptional specialization post-fusion we observed here. From these observations, we posit that the same microtubule dynamics known to promote myonuclear transcriptional specialization toward NMJ transcriptional signature is a general MuSC-Derived myonuclear adaptation that also applies to MTJ transcriptional specialization we observed in Adult. We attribute the divergent post-fusion differentiation of these MuSC-Derived myonuclei to sensing denervation stress, as we only identified denervated myonuclei in aged muscle, consistent with previous reports of progressive denervation with advancing age^34–36^. Resident myonuclei also appear to have the capacity to support NMJ myonuclei during MOV with Adult resident myonuclei having a greater magnitude of response than Aged muscle^8^. MuSC-Derived myonuclei specializing toward an NMJ myonuclear transcriptional signature in Aged muscle then may be compensating for impaired responsiveness of resident myonuclei. Lastly, we observed age-dependent expression of *Nav3* in MTJ and NMJ clusters in Adult and Aged mice, respectively. We also observed modest upregulation of *Nav3* transcript abundance in Transitioning myonuclei, suggesting *Nav3* may play a role in physically guiding myonuclei to their terminal location.

We next sought to determine transcriptional effectors of MuSC-Derived myonuclear specialization. Elevated *Runx1* expression in transitioning MuSC-Derived myonuclei provided evidence *Runx1* may be involved MuSC-Derived myonuclear differentiation. Testing this in-silico, we found evidence that *Runx1* may be a critical regulator of the transition state of MuSC-Derived myonuclei, independent of age. This is consistent with our previous finding where *Runx1* expression was higher in Transitioning myonuclei, but nearly absent in terminal state clusters (MTJ and Spindle in Adult, NMJ and Spindle in Aged). In addition, we found in Adult muscle, *Runx1* expression was almost exclusively expressed by MuSC-Derived myonuclei, but in Aged muscle it was expressed in MuSC-Derived and Denervated myonuclei, suggesting potential roles of Runx1 in both pathology and load-induced adaptation^8,30^.

Aside from MuSC-Derived myonuclear specialization in typical (extrafusal) fibers, we found that MuSC-Derived myonuclei in Spindle (intrafusal) fibers have a unique transcriptional trajectory post-fusion. Interestingly, intrafusal fibers are more dependent on MuSCs to maintain their size and in-vivo functional output in both Adult^37^ and Aged^38^ muscle. Despite this being the sole functional output of MuSC depletion, molecular interrogation of this phenomenon is completely absent in existing literature. Understanding the role of MuSC-Derived myonuclei in intrafusal fibers will position these MuSC-Derived myonuclei as a therapeutic target for aging, where age-associated loss of coordination can lead to a fatal fall^39^. To determine this, we distinguished these intrafusal MuSC-Derived myonuclei using a computed likelihood of transcriptional trajectory into the Spindle myonuclear cluster. A defining feature that distinguished intrafusal from extrafusal MuSC-Derived myonuclei was markedly higher expression of *Esrrg*, a transcription factor known to associate with aerobic adaptations and elevated expression of oxidative metabolism gene programs^26,27^. Resident spindle myonuclei are dominated by expression of an oxidative program including expression of Type I myonuclear genes (*Myh7*, *Atp2a2*, *Tpm3*). Our *in-silico* transcription factor KO screen showed Esrrg is a driving transcription factor of MuSC-Derived myonuclear differentiation in intrafusal fibers only. These data suggest Esrrg may promote the mature intrafusal fiber myonuclear transcriptional program.

In conclusion, we found 1) the transcriptional signature of MuSC-Derived and resident MOV-Responsive myonuclei, 2) MuSC presence is important for Aged but not Adult myonuclear transcriptional responsiveness to MOV, 3) a subset of MuSC-Derived myonuclei that are transitioning toward a more specialized state, marked by increased expression of *Runx1* and 4) that MuSC-Derived myonuclei behave differently in Adult and Aged muscle, but in both cases, adopt highly specialized transcriptional signatures that are distinct from fiber type-specific myonuclei, and that *Nav3* and microtubule remodeling likely play a role in such specialization. Ultimately, we posit that MuSC-Derived myonuclei are critical for maintaining specialized structures in skeletal muscle, potentially explaining a lack of MuSC dependence for maintenance and adaptation of muscle size in Adult and Aged muscle^5,6^.

## Methods

### Mouse model and mechanical overload surgery

All animal studies were performed in accordance with institutional guidelines and approved by the Institutional Animal Care and Use Committee of the University of Kentucky. Mice were housed in a temperature-and humidity-controlled room, maintained on a 14:10-h light-dark cycle, and food and water were provided ad libitum throughout the entire study. Mice used in this study had two genetic strategies 1) the myofiber-specific Tet-On system to label myonuclei as previously described for the HSA-rtTA^+/−^;TRE-H2B-GFP^+/−^ (HSA-GFP) mice^40^ and 2) the Pax7^Cre/+^;Rosa26^DTA/+^ (Pax7-DTA) model that allows tamoxifen satellite cell depletion as previously described^6^. Mice (female, Adult = 6 months, Aged = 24 months of age) were treated with tamoxifen-free vehicle (15% ethanol in sunflower seed oil) or tamoxifen (2 mg/day) for five consecutive days. Following a 14-day washout period, all mice underwent either surgically induced synergist ablation surgery to induce mechanical overload (MOV) of the plantaris muscle as previously described^41^, or Sham surgery. After 9 days of MOV, mice were given doxycycline (0.5 mg/mL, 2% sucrose) for 5 days for a total of 14 days of MOV prior to tissue collection. This strategy leads to the labeling of approximately 90%-95% of myonuclei, with negligible off-target labeling of nonmyonuclei^40^, allowing fluorescence activated sorting of myonuclei, specifically.

### Single-nucleus RNA sequencing

Raw sequencing reads were aligned to the mm39 reference genome and quantified using Cell Ranger, producing per-sample filtered feature-barcode matrices that were used as input for all downstream analyses. Per-sample quality control and downstream analysis was performed using Scanpy (v1.12). Nuclei were required to express a minimum of 200 genes. Upper and lower gene-count boundaries were set at the 99th and 1st percentiles of each sample’s gene-count distribution to remove outlier nuclei. Nuclei in which mitochondrial or hemoglobin transcripts exceeded 5% of total counts were excluded as low-quality or contamination. Genes detected in fewer than 10 cells within a sample were discarded prior to sample-level filtering. Following per-sample filtering, all samples were concatenated and a second round of gene filtering (minimum 10 cells across the combined dataset) was applied. Doublets were identified per sample using a two-stage approach. A variational autoencoder was first trained independently on each sample with scVI (v1.4.2) (10 latent dimensions), and the resulting latent representation was used to train a SOLO binary classifier. Nuclei with a SOLO doublet score exceeding 0.15 were removed. Raw counts were preserved in a dedicated layer (counts) for integration. Library sizes were normalized to 10,000 counts per nucleus followed by log(1+x) transformation. Highly variable genes (HVGs) were selected in a batch-aware manner (stratified by sample) using the Seurat v3 flavor, retaining the top 3,000 HVGs for integration. Batch effects across samples were corrected using scVI. The scVI model was trained on raw counts with a 10-dimensional latent space. The scVI latent representation was used to construct a shared nearest-neighbor graph (k = 15 neighbors, 50 PCs) for UMAP embedding and Leiden clustering (v0.11.0) (resolution = 1.0). Cluster identities were assigned by scoring each cluster against a curated consensus marker gene set drawn from McKellar et al. (2021)^42^, PanglaoDB, and CellMarker, using scanpy.tl.score_genes. Differential expression between clusters was performed with the Wilcoxon rank-sum test in Scanpy. Nuclei annotated as myofiber subtypes were extracted from the full dataset and subjected to independent reclustering. HVG selection was rerun retaining 9,000 genes (Seurat flavor), followed by PCA (50 components), nearest-neighbor graph construction (k = 10), and Leiden clustering (resolution = 1.0). UMAP was computed with min_dist = 0.1 and spread = 2.0 to improve separation of transcriptionally similar fiber-type subtypes. After preprocessing and quality control, we had a total of 21,255 myonuclei across the full dataset (Adult Sham MuSC**^+^** = 2,277; Adult MOV MuSC**^+^** = 2,445; Adult Sham MuSC− = 2,322; Adult MOV MuSC− = 2,027; Aged Sham MuSC**^+^**= 2,969; Aged MOV MuSC**^+^** = 2,274; Aged Sham MuSC− = 3,130; Aged MOV MuSC− = 3,811). Wilcoxon rank-sum differential expression was performed across reclustered myonuclear populations to identify cluster-specific marker genes.

### Kernel Density Estimate and differential density analysis

Two-dimensional kernel density estimates (KDE) were computed for each experimental condition (MuSC**^+^**, MuSC**^-^**, Sham, MOV, Adult, Aged) and each individual sample independently using the gaussian_kde function from scipy.stats (scipy v1.16.3), and applied to each nucleus’ UMAP embedding coordinates. The kernel bandwidth was selected automatically using the default Scott’s rule, which adapts the degree of smoothing to the number of nuclei in each density estimate. For visualization and differential analysis, each pairwise KDE comparison was evaluated on an equal grid spanning the full UMAP. Condition-specific densities were normalized to the same scale by dividing by the maximum value across grids for each sample, yielding densities values 0-1. Differential density was computed as the difference between the two normalized grid surfaces and interpolated back to individual cell positions using RegularGridInterpolator with linear interpolation. Per-nucleus density values were computed by evaluating each condition’s density measurement directly at the UMAP coordinates in which the nucleus resides. Spatially coherent regions of differential density enrichment were identified using DBSCAN (Density-Based Spatial Clustering of Applications with Noise; scikit-learn v1.8.0) applied to the UMAP coordinates of cells meeting a differential density threshold < 0.1. Clustering was performed with sklearn.cluster.DBSCAN using a neighborhood radius of eps = 0.8 and a minimum cluster size of 30 nuclei. For each identified cluster, a convex hull was computed using scipy.spatial.ConvexHull to define the spatial boundary of the population. The largest cluster (by nucleus count) was retained as the primary region of interest for downstream analyses, and only cells whose UMAP coordinates fell within the corresponding convex hull were assigned membership to the DBSCAN-identified cluster.

### Lasso regression and differential density prediction

To identify genes whose expression predicts spatial enrichment within the MuSC-Derived myonucleus population, regularized regression was performed within the MuSC-Derived convex hull. Raw count profiles for hull-contained nuclei were used as differential density predictors and all gene features were standardized to zero mean and unit variance using sklearn.preprocessing.StandardScaler. A LassoCV (scikit-learn v1.8.0) model was fit with 5-fold cross-validation to optimize the regularization parameter, α over a maximum of 10,000 iterations. The cross-validation–selected α was used to refit a final Lasso model, and the predicted differential density was evaluated against observed values using spearman correlation coefficient, r, and R². Genes with non-zero Lasso coefficients comprised the gene set used in the final model to predict differential density with positive lasso coefficient predicting high differential density and negative lasso coefficients predicting low differential density.

### Trajectory analysis

Following normalization (sc.pp.normalize_total, sc.pp.log1p), PCA, nearest-neighbor graph construction, and UMAP recomputation, a diffusion map was computed using scanpy.tl.diffmap. The trajectory root was defined as the MuSC-Derived nucleus closest to the centroid of the MuSC-Derived cluster in UMAP space. Diffusion pseudotime (DPT) was then computed with scanpy.tl.dpt anchored to this root nucleus. A pseudotime-based transition matrix was computed using CellRank’s (v2.2.0) PseudotimeKernel^19^. Terminal cell states were estimated using the Generalized Perron Cluster Cluster Analysis (GPCCA) estimator implemented in CellRank (cellrank.estimators.GPCCA), initialized with the pseudotime-based transition matrix. Terminal states were predicted automatically and fate probabilities toward each terminal state were computed with g.compute_fate_probabilities. To quantify the rate of fate commitment across the UMAP landscape, a local gradient of fate probability was computed in PCA space. For each nucleus, the 30 nearest neighbors by Euclidean distance in PCA space were identified using a k-d tree (scipy.spatial.cKDTree) constructed on the first 20 principal components. The gradient magnitude at each cell was defined as the mean Euclidean distance in fate-probability space between that cell and each of its neighbors, normalized by the corresponding distance in PCA space.

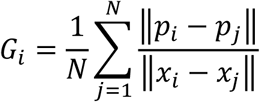

Where G_i_ is the fate commitment gradient magnitude for nucleus i, N = 30 nearest neighbors of nucleus i in PCA space, p_i_ and p_j_ are the fate probability vectors (across all terminal states) for nucleus i and its neighbor j, and x_i_ and x_j_ are the corresponding coordinates in the first 20 principal components.

### In-silico transcription factor knockout screening

To predict the effect of individual transcription factor (TF) loss-of-function on myonuclear differentiation trajectories, we performed in-silico knockout (KO) simulations using CellOracle (v0.20.0) on age-matched (Adult, Aged) gene regulatory network (GRN) models fit to the vehicle-treated single-myonucleus RNA-seq dataset using a base GRN built from CellOracle’s pre-packaged mouse scATAC-seq-atlas promoter-accessibility reference. Gene expression was imputed by k-nearest-neighbor smoothing following PCA, with the number of principal components selected by an elbow heuristic on the explained-variance curve (clamped to 10–50 components) and neighborhood size set to 2.5% of the total nucleus count per age (minimum 15 neighbors, balanced KNN). Cluster-specific GRN edges were fit by Bayesian ridge regression (grouped by annotated cell type, regularization α = 10) and filtered to the top 2,000 edges per network by absolute coefficient weight at p < 0.001, after which per-cluster network centrality statistics (degree, betweenness, and eigenvector centrality) were computed for downstream regulator selection. Candidate TFs were selected as hub regulators of the MuSC-Derived subnetwork by GRN out-degree/eigenvector centrality, restricted to those with adequate baseline expression (≥5% of nuclei expressing).

For each candidate TF, expression was set to zero across all nuclei (simulate_shift, perturb_condition=TF: 0.0, n_propagation=3), and the predicted post-perturbation cell state transition probabilities were estimated by KNN-based Markov transition modeling (estimate_transition_prob, n_neighbors=200, knn_random=True, sampled_fraction=1) and projected onto the UMAP embedding as a per-nucleus displacement vector (calculate_embedding_shift, sigma_corr=0.05).

To quantify whether a given KO shift promoted or opposed precomputed differentiation, we computed an inner-product score between the simulated shift and a reference differentiation-flow field, defined as the local gradient of diffusion pseudotime (dpt_pseudotime) across the embedding (CellOracle Gradient_calculator, mass-filtered at a minimum local cell density of 0.01). The reference field and its gradient were computed once per age and shared across all TF perturbations. Inner products were calculated on CellOracle’s coarse representative grid (Oracle_development_module.calculate_inner_product), with grid points falling below the local-mass threshold excluded from downstream analysis. By convention, a negative inner product indicates the KO shift opposes normal differentiation progression (i.e., the TF normally promotes/drives that transition), while a positive value indicates the shift is aligned with normal progression (i.e., the TF normally restrains/represses it). A matched randomized-control inner product was computed natively by CellOracle for each grid point to assess the null distribution of shift-flow alignment expected in the absence of a directional perturbation effect.

For terminal state restricted analyses, each grid point was assigned the identity of its nearest real nucleus in UMAP space (nearest-neighbor search, scipy cKDTree), yielding both a cluster identity (whether the nucleus belongs to the corresponding terminal population) and, for MuSC-Derived myonuclei, a predicted terminal stat-target label. Terminal state-target labels were derived by z-scoring each MuSC-Derived myonucleus’ CellRank (GPCCA) absorption probability toward each terminal state and assigning the nucleus to the terminal state with the maximum z-scored probability, correcting for differences in baseline absorption-probability magnitude across terminal states. The terminal state-specific TF screen value was then defined as the mean of the general (dpt_pseudotime-referenced) inner product described above, restricted to grid points whose nearest nucleus was either predicted to be terminal state-targeted, or a member of the corresponding terminal cluster itself i.e., pooling nuclei along the full trajectory from lineage commitment to terminal differentiation, rather than the terminal-state targeted alone to capture the full differentiation trajectory including both the transitioning MuSC-Derived myonuclei and the terminal state which those myonuclei are targeting.

## Acknowledgements

The authors thank the personnel at the University of Kentucky Genomics Core Laboratory for assistance with sequencing experiments, the University of Kentucky Flow Cytometry and Immune Monitoring Core for technical support with cell sorting, and Dr. Doug Harrison of the University of Kentucky Biology Department/Genetics for support with 10X Genomics. This work was supported by NIH grant R01 AG069909 to JJM, CAP, and YW.

## Authors’ Contributions

JZG and NTT performed experiments and collected data. JZG, and AI conducted animal surgeries and tissue processing, performed flow cytometry and cell sorting. NTT and YW performed bioinformatics analyses. AI, YW, CAP, and JJM conceived the study, designed experiments, and provided supervision. KAM provided intellectual contributions and expertise for experimental design and manuscript preparation. JJM, YW, CAP, and CSF provided resources, intellectual contributions, and expertise. NTT and YW prepared figures and drafted the manuscript. All authors reviewed, edited, and approved the final manuscript.

## Figure Legends

<u>*Figure S1: Lasso regression performance for MuSC-Derived and MOV-Responsive clusters.*</u>

A. Reconstruction of the 3D differential density shape from Lasso regression-based feature selection which produced a 323 gene model. B. Quantification of Lasso regression performance predicting differential density with the 323 gene model. C. Relationship between Lasso coefficient and pearson correlation between log1p expression and distance to differential density peak for genes in the 323 gene model. D. Expression pattern of genes with positive Lasso coefficients. E. Lasso regression performance in the MOV-Responsive cluster. F. Distribution of differential density across MuSC-Derived and MOV-Responsive myonuclear clusters.

## Conflict of Interest

YW is the founder of MyoAnalytics LLC. AI, JJM, and YW are co-founders of Myobiota, Inc.

