## Supplementary figures and images for "Skeletal Muscle Stem Cell-Derived Myonuclei Adopt Divergent Terminal Transcriptional States in Adult and Aged Muscle In Response to a Hypertrophic Stimulus"

### Supplemental Figure 1

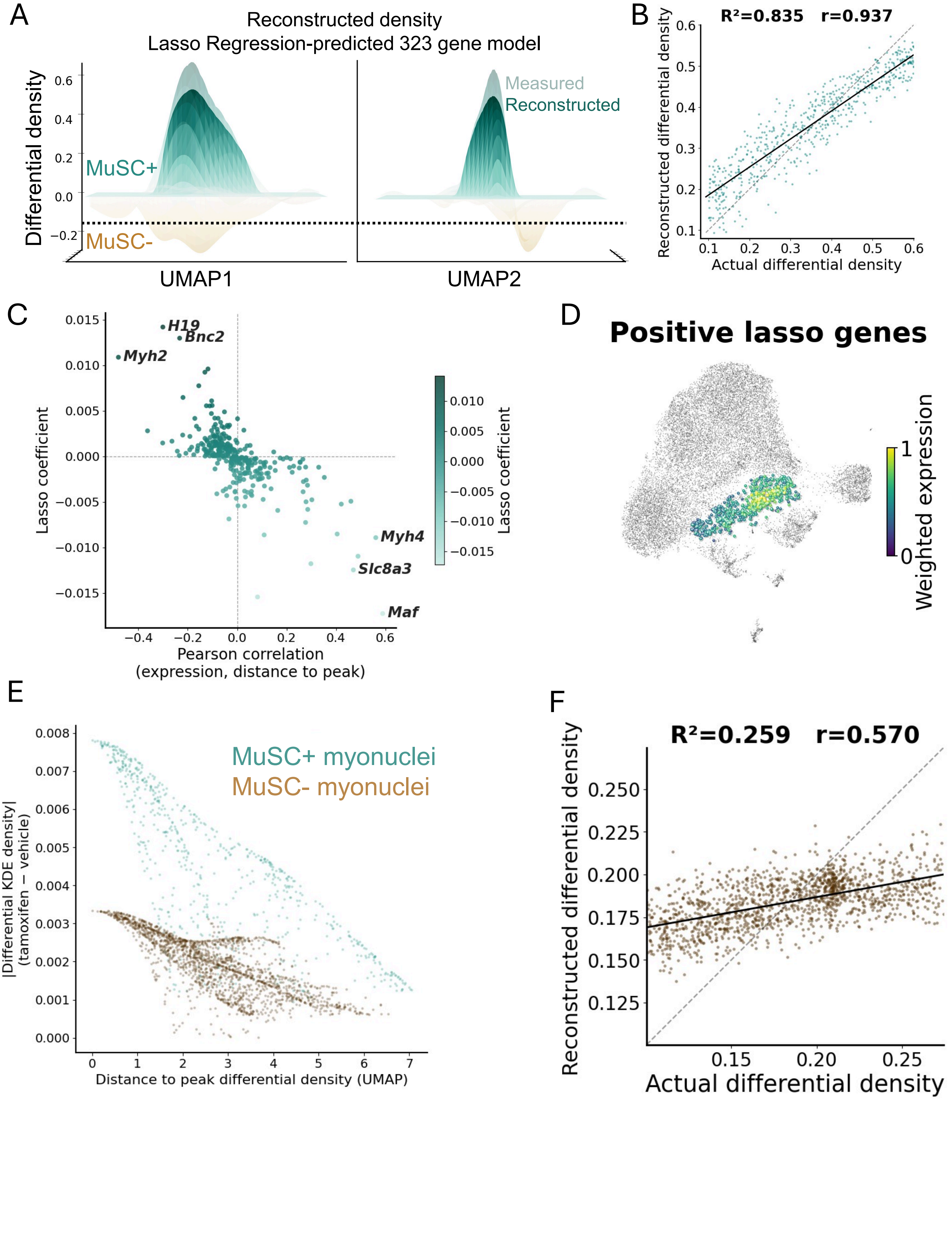
